# The evolution of two transmissible leukaemias colonizing the coasts of Europe

**DOI:** 10.1101/2022.08.06.503021

**Authors:** Alicia L. Bruzos, Martín Santamarina, Daniel García-Souto, Seila Díaz, Sara Rocha, Jorge Zamora, Yunah Lee, Alejandro Viña-Feás, Michael A. Quail, Iago Otero, Ana Pequeño-Valtierra, Javier Temes, Jorge Rodriguez-Castro, Antonio Villanueva, Damián Costas, Rosana Rodríguez, Tamara Prieto, Laura Tomás, Pilar Alvariño, Juana Alonso, Asunción Cao, David Iglesias, María J. Carballal, Ana M. Amaral, Pablo Balseiro, Ricardo Calado, Bouchra El Khalfi, Urtzi Izagirre, Xavier de Montaudouin, Nicolas G. Pade, Ian Probert, Fernando Ricardo, Pamela Ruiz, Maria Skazina, Katarzyna Smolarz, Juan J. Pasantes, Antonio Villalba, Zemin Ning, Young Seok Ju, David Posada, Jonas Demeulemeester, Adrian Baez-Ortega, Jose M. C. Tubio

## Abstract

Transmissible cancers are malignant cell clones that spread among individuals through transfer of living cancer cells. Several such cancers, collectively known as bivalve transmissible neoplasia (BTN), are known to infect and cause leukaemia in marine bivalve molluscs. This is the case of BTN clones affecting the common cockle, *Cerastoderma edule*, which inhabits the Atlantic coasts of Europe and north-west Africa. To investigate the origin and evolution of contagious cancers in common cockles, we collected 6,854 *C. edule* specimens and diagnosed 390 cases of BTN. We then generated a reference genome for the species and assessed genomic variation in the genomes of 61 BTN tumours. Analysis of tumour-specific variants confirmed the existence of two cockle BTN lineages with independent clonal origins, and gene expression patterns supported their status as haemocyte-derived marine leukaemias. Examination of mitochondrial DNA sequences revealed several mitochondrial capture events in BTN, as well as co-infection of cockles by different tumour lineages. Mutational analyses identified two lineage-specific mutational signatures, one of which resembles a signature associated with DNA alkylation. Karyotypic and copy number analyses uncovered genomes marked by pervasive instability and polyploidy. Whole-genome duplication, amplification of oncogenes *CCND3* and *MDM2*, and deletion of the DNA alkylation repair gene *MGMT*, are likely drivers of BTN evolution. Characterization of satellite DNA identified elements with vast expansions in the cockle germ line, yet absent from BTN tumours, suggesting ancient clonal origins. Our study illuminates the evolution of contagious cancers under the sea, and reveals long-term tolerance of extreme instability in neoplastic genomes.

## Introduction

Transmissible cancers are clonal somatic cell lineages that spread between individuals via direct transfer of living tumour cells, in a process analogous to cancer metastasis (*1, 2*). Naturally occurring transmissible cancers have been identified in dogs (*3–5*), Tasmanian devils (*6–8*) and, more recently, several species of marine bivalve molluscs (*9–14*). To date, eight transmissible cancer lineages, collectively known as bivalve transmissible neoplasia (BTN), have been described in bivalves, probably spreading via transfer of free-floating cells in seawater. BTN infection causes a leukaemia-like disease termed disseminated neoplasia (DN), where neoplastic cells proliferate and accumulate in the host’s haemolymph and solid tissues (*15*). Although DN is generally fatal, slow progression and remission have been described (*16, 17*).

Among the species affected by DN is the common cockle, *Cerastoderma edule*. This marine bivalve is distributed along the Atlantic coasts of Europe and north-west Africa, being typically found in tidal flats at bays and estuaries (*18*). Adult cockles bury themselves in the seabed sediment and use their syphons and gills to filter seawater for sustenance. DN in common cockles was first documented 40 years ago in Ireland (*19*), and later identified in other European countries (*15*). A genetic study recently provided evidence that DN in *C. edule* is caused by transmissible cancer, and suggested the existence of at least two BTN clones in this species (*10*). Nevertheless, the origins and evolution of cockle BTN lineages remain entirely unexplored.

Here, we present the first comprehensive study of the genomes of BTN clones affecting *C. edule* in Europe. We sampled thousands of common cockle specimens across 11 countries, obtained a chromosome-level reference genome for the species, and used it to catalogue the genomic variation in 61 BTN tumours identified in these animals. Combining histopathology, cytogenetics, and sequencing of whole genomes and transcriptomes, our study illuminates the evolutionary history of the marine leukaemias that have colonized European cockle populations.

### Prevalence of disseminated neoplasia in common cockles

To investigate the current prevalence of DN in *C. edule*, we collected 6,854 specimens at 36 locations from 11 countries along the Atlantic coasts of Europe and north Africa between 2016 and 2021 (**Fig. 1A; Table S1**). This included intensive sampling on the coasts of Ireland and Galicia (north-west Spain), two regions where high prevalence of DN has been reported in the past (*20–22*). Cytohistological examination of haemolymph and solid tissues revealed that 5.7% (390/6,854) of specimens were infected by abnormal circulating cells displaying the features of DN (**Fig. 1A; Table S1**). High overall prevalence was observed in Portugal (17.6%), Ireland (7.4%) and Spain (6.4%), with lower prevalence found in the United Kingdom (3.6%) and France (1.1%; **Fig. S1**); no DN cases were detected in the remaining six countries (Germany, Denmark, Morocco, the Netherlands, Norway, Russia). Twenty percent (77/390) of neoplastic specimens presented a severe form of the disease (stage N3), characterized by high levels (>75%) of neoplastic cells in the haemolymph and massive tissue infiltration foci; 26% (102/390) presented an intermediate form (stage N2), distinguished by 15–75% of neoplastic cells in the haemolymph and presence of small infiltration foci in one or more organs; the remaining individuals (53%, 208/390) were diagnosed with a mild form (stage N1), where low levels (<15%) of neoplastic cells circulate in the haemolymph and infiltrate solid tissues in small numbers (*22*) (**Fig. S2; Table S2**).

**Figure 1.**
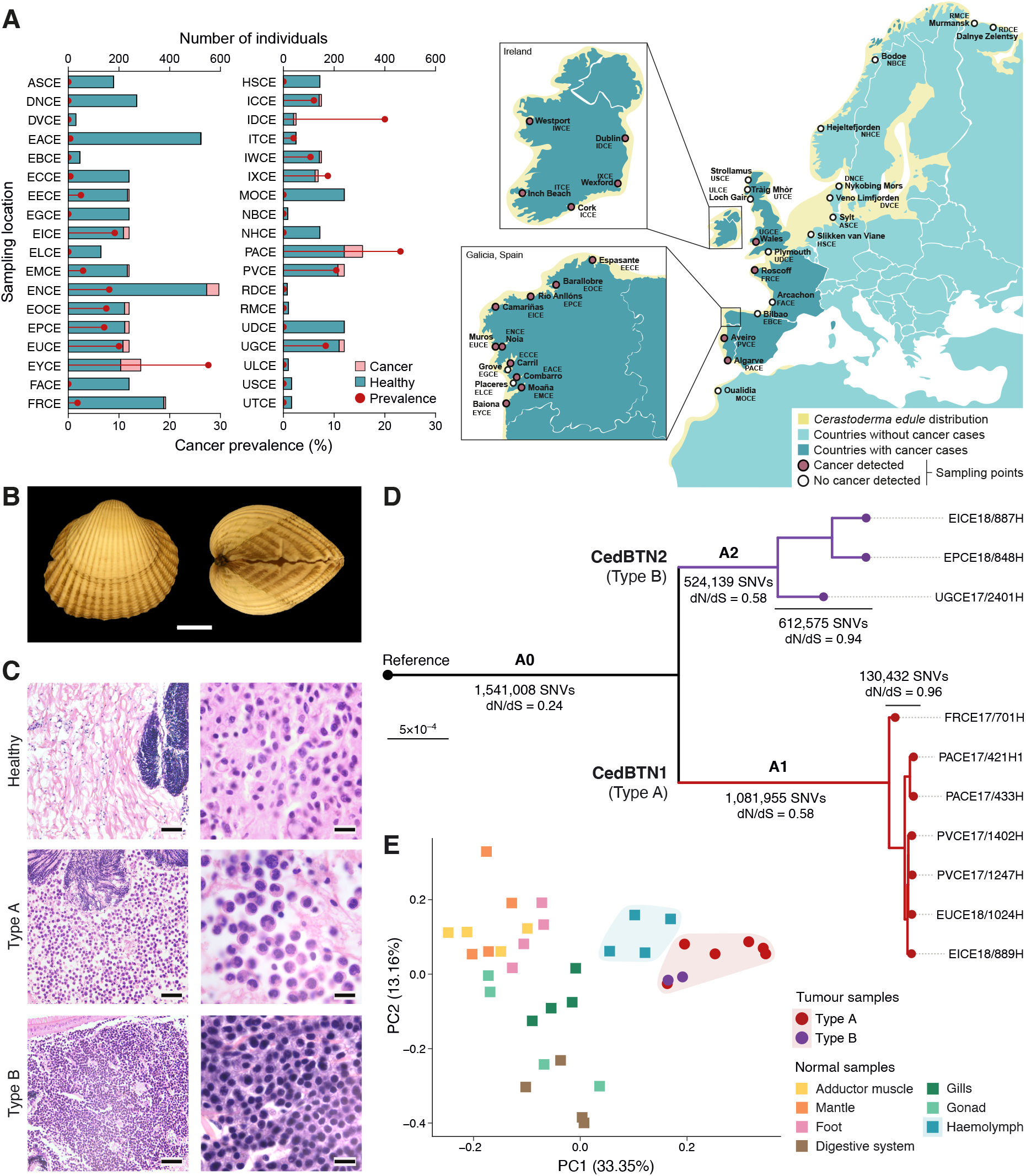
Distribution, origins and clonal structure of transmissible neoplasia in common cockles. **A**, Numbers of healthy and neoplastic *C. edule* cockles collected at each sampling location, with overall cancer prevalence per location for 2016–2021 (left). Map shows sampling locations and geographic distribution of the species. **B**, Photographs of the individual from which the reference *C. edule* genome was assembled (scale bar, 10 mm). **C**, Micrographs of histological sections from healthy and DN-affected cockle tissues. Images in the lefthand column show healthy connective tissue surrounding the male gonadal follicle (top) and connective tissue heavily infiltrated by type A and type B DN cells (scale bars, 50 μm). Images in the right-hand column show details of normal haemocytes (top), type A and type B DN cells (scale bar, 10 μm). **D**, Phylogenetic tree inferred from BTN-specific SNVs in 10 high-purity tumour samples, showing concordance between histological DN types A and B and two clonal transmissible cancer lineages, CedBTN1 and CedBTN2. Numbers of SNVs and dN/dS ratios are provided for different sections of the tree. All nodes have bootstrap support values of 100 (*n*=1000 replicates). Scale bar indicates phylogenetic distance (SNVs per site). **E**, Principal component analysis of gene expression for genes with tissue-specific expression in normal cockle tissues and DN samples, indicating a clustering of DN (red shading) with healthy haemolymph (blue shading).

### Reference genome and transcriptome of the common cockle

As an initial step in our genomic study of cockle DN, we applied multiplatform DNA sequencing to obtain a reference assembly of the *C. edule* genome (**Fig. S3**). As our reference specimen, we selected a healthy adult male cockle (**Fig. 1B**) carrying a standard karyotype with 19 chromosome pairs. Hybrid genome assembly yielded a chromosome-level reconstruction of the cockle nuclear genome into 19 scaffolds (N50=39.6 megabases, Mb; **Table S3**), with an additional 14.9-kilobase (kb) scaffold containing the mitochondrial genome sequence. Haploid genome size was estimated at 790 Mb, with a G+C content of 35.6%. We additionally employed RNA sequencing data from seven tissues to reconstruct a 290-Mb reference transcriptome displaying 98.8% completeness in metazoan gene content (**Table S3**). Gene annotation resulted in a 42-Mb exome with 14,055 protein-coding genes. While this protein-coding exome constitutes 5.3% of the total nuclear genome size, repetitive sequences comprise 46.2% of the genome, with long interspersed nuclear elements (LINEs) being the most frequent type of transposable element among annotated repeats (**Fig. S4; Table S4**).

### Two transmissible cancer lineages propagate through cockle populations

Traditionally, two distinct classes of cockle DN, termed types ‘A’ and ‘B’, have been described through cytohistological methods, on the basis of differences in tumour cell size and morphology (*21*) (**Fig. 1C**). A previous analysis of microsatellite variation and single-nucleotide variants (SNVs) in both mitochondrial DNA (mtDNA) and one nuclear gene provided evidence that these DN types represent two transmissible cancer lineages (*10*), although it is possible that further lineages, as well as non-transmissible cases of cockle DN, exist.

To further investigate the origins and evolution of cockle BTN, we performed whole-genome sequencing of neoplastic haemolymph samples from 61 individuals diagnosed with DN (**Table S5**). Ten of these samples, presenting high (≥90%) tumour purity, were designated as a BTN ‘golden set’, and used to identify a collection of high-confidence candidate somatic variants. We also sequenced matched tissue samples from 40 host individuals and 462 non-neoplastic specimens collected across the species’ geographic range (**Table S5**). After accounting for host DNA contamination and common germline polymorphisms, we identified a total of 4.3 million SNVs (2.5–3.1 million SNVs per sample; **Table S6**) and 0.7 million short insertions and deletions (indels). This ‘BTN-specific’ variant set includes both somatic variants in each BTN lineage and ancestral germline polymorphisms (from each lineage’s founder individual) which were absent from our panel of 462 non-neoplastic cockles.

We used BTN-specific SNVs to reconstruct a tumour phylogenetic tree, which split the 10 ‘golden set’ tumours into two divergent lineages (**Fig. 1D**), consistently corresponding to the two histological types of cockle DN (**Fig. S5**). We hereafter refer to these two lineages of *C. edule* BTN, respectively corresponding to DN types A and B, as CedBTN1 and CedBTN2. To assess the quality of our variant set and confirm the independent origins of both BTN lineages, we estimated the ratio of nonsynonymous to synonymous mutation rates (dN/dS) (*23, 24*) along the phylogenetic tree (**Fig. 1D**). The dN/dS ratios for variants shared by all 10 tumours (ancestral variant set ‘A0’) and variants shared by all tumours in each lineage (sets ‘A1’ and ‘A2’) indicate that these sets contain a large fraction of germline polymorphisms from two separate founder individuals (dN/dS 0.24 for A0, 0.58 for A1, 0.58 for A2). In contrast, the dN/dS for the terminal branches approximates a neutral value of 1.0 (0.96 for CedBTN1, 0.94 for CedBTN2), as expected for pure sets of somatic mutations (*24*). Accordingly, the dN/dS of variants found in only one tumour (private variants) is 1.00 (**Table S7**).

Additionally, we performed principal component analysis on a set of germline polymorphisms genotyped across the 10 ‘golden set’ tumours and 100 non-neoplastic cockles covering all sampled populations (**Fig. S6**). This analysis split the tumours into two divergent clusters matching CedBTN1 and CedBTN2, and set apart from two non-neoplastic sample clusters representing relatively divergent groups of cockle populations from northern and southern Europe (*25*). This result suggests that CedBTN clones are highly divergent both from each other and from modern cockle populations, further supporting two independent origins. Nevertheless, analysis of sequence mapping data shows that the fractions of sequence reads aligning against the *C. edule* reference genome in BTN samples (97.4–98.1%, ‘golden set’ samples) are comparable to those for 462 non-neoplastic cockles (interquartile range 97.1–97.8%) and substantially higher than fractions for cockles of the closest known species, *C. glaucum* (48.3–60.4%, six samples), consistent with both clones having arisen in *C. edule* animals.

### Haemocytic origin of cockle transmissible neoplasia

The ontogeny of bivalve DN is a long-standing question with relevance for the biology and evolution of BTN. The fact that DN cells are observed in the circulatory system and share morphological features with haemocytes has traditionally led to their consideration as neoplastic haemocytes (*15*). Nevertheless, some studies have proposed alternative tissues of origin for these cancers, including the gonad follicles or gill epithelium (*15*).

To shed light on the origins of CedBTN lineages, we sequenced the transcriptomes of haemolymph samples from eight cockles diagnosed with advanced stages DN, and a collection of seven tissues (adductor muscle, mantle, foot, digestive system, gills, gonad and haemolymph) from 28 non-neoplastic animals (**Table S5**). Gene expression analysis for a set of 420 genes with tissue-specific expression (60 genes per tissue type) indicated a consistent gene expression profile for type A and type B DN samples, which is close to that of non-neoplastic haemolymph samples and divergent from those of all other tissues (**Fig. 1D; Fig. S7; Table S8**). This finding suggests that cockle BTN clones are cancers of the haemolymphatic system, derived from somatic haemocytes or haemic progenitor cells. This recurrent cellular origin may reflect an exclusive capability of malignant haemocytes to exploit the transmission opportunities offered by the open circulatory system of bivalves.

### Mitochondrial transfer delineates the clonal structure of CedBTN

To explore the evolutionary history of CedBTN at the mitochondrial level, we identified SNVs in the mtDNA of 51 haemolymph samples from neoplastic cockles, 40 matched-host tissue samples and 168 non-neoplastic cockle samples. In neoplastic animals, sequencing data showed two mtDNA haplotypes at distinct variant allele fractions (VAFs), corresponding to the host and CedBTN mitochondrial genomes. Combining tumour purity and mtDNA VAF information to deconvolute the mtDNA haplotypes within each sample, we identified nine distinct tumour haplotypes (six in CedBTN1 and three in CedBTN2), each distinguished by a specific set of mtDNA variants (**Fig. 2A; Table S6; Table S9**).

**Figure 2.**
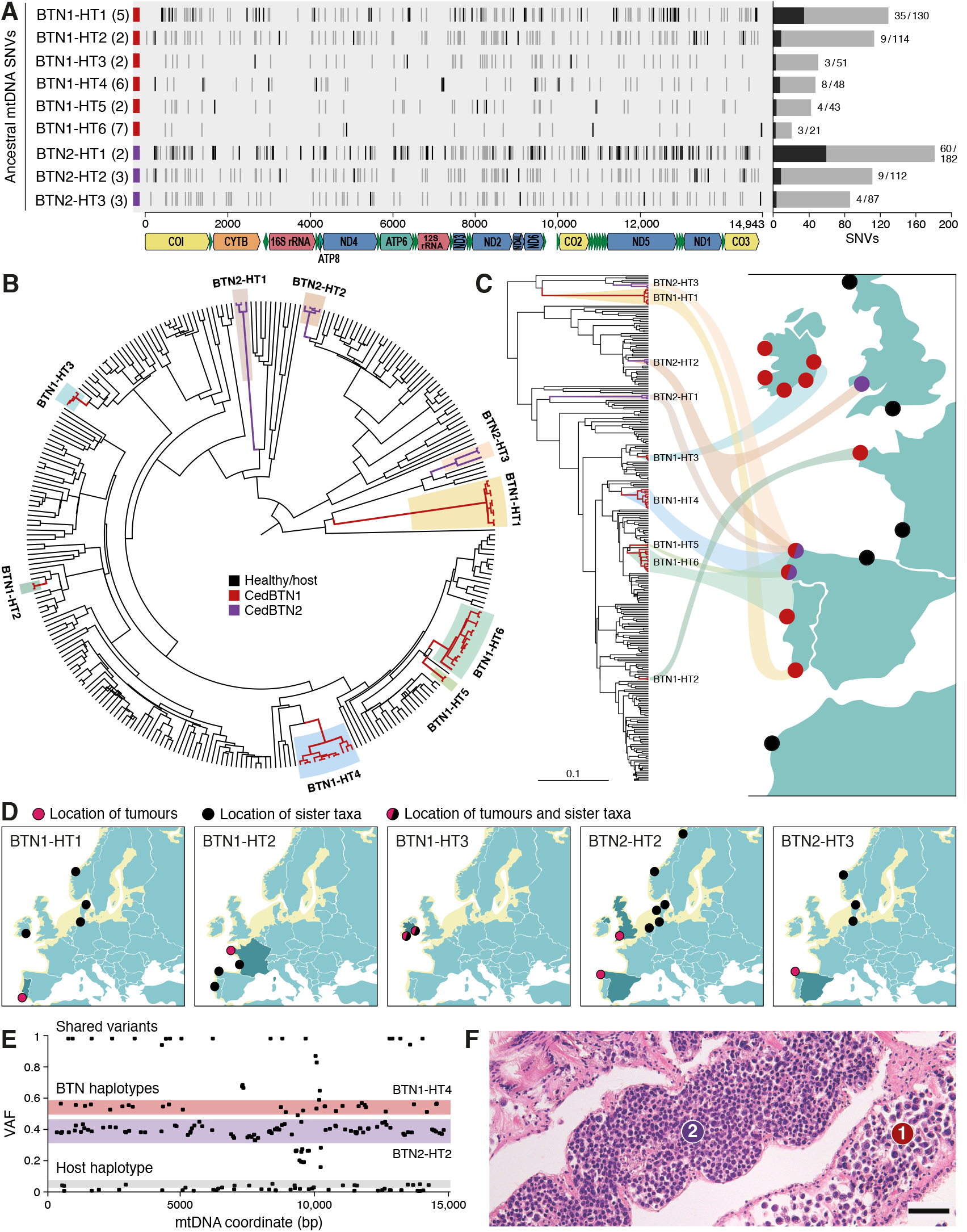
Mitochondrial DNA phylogeny, mtDNA horizontal transfer and co-infection in CedBTN. **A**, Ancestral mtDNA haplotypes identified in CedBTN samples, with ancestral SNVs (common to all samples carrying the haplotype) arranged along the reference mtDNA sequence (*x*-axis). Potentially somatic SNVs (absent from non-neoplastic samples) are shown in black. Potential mtDNA HT events associated with each haplotype in CedBTN1 (red) and CedBTN2 (purple) are labelled, with number of samples used to identify ancestral variants given in parentheses. Bar plot presents numbers of potentially somatic (black) and total (grey) ancestral variants per haplotype; numbers are indicated next to each bar. A schematic representation of the mtDNA gene annotation is shown at the bottom. **B**, Bayesian phylogenetic tree of mtDNA haplotypes in normal and CedBTN samples, with identified tumour mtDNA lineages highlighted and labelled. Branch lengths represent phylogenetic distance (scale bar given in **C**). **C**, Correspondence between mtDNA phylogenetic tree and tumour sampling regions; map point colours denote CedBTN lineages as in **B**. Sampling points in Galicia (north-west Spain) are grouped into northern and southern points. Scale bar indicates phylogenetic distance (SNVs per site). **D**, Maps showing locations of tumours and normal sister taxa for five mtDNA lineages. **E**, VAF plot evidencing co-infection of a host (EICE18/910) by cells from two mtDNA lineages, one from each CedBTN clone. Three observed mtDNA haplotypes are shaded in different colours. **F**, Micrograph of histological section of gills from EICE18/910, confirming co-infection by both CedBTN clones. Dilated efferent vessels are shown; vessels labelled ‘1’ and ‘2’ are mainly infiltrated by type A and type B neoplastic cells, respectively. Scale bar, 50 μm.

The findings above suggested the existence of nine CedBTN mtDNA lineages. This was confirmed through phylogenetic reconstruction via maximum likelihood (ML) and Bayesian methods (**Fig. 2B; Fig. S8; Fig. S9**). The presence of multiple mtDNA lineages within each CedBTN clone indicates that mitochondria from transient hosts have repeatedly been acquired by these tumours, as previously described for other transmissible cancers (*26, 27*). We therefore labelled these mtDNA lineages after putative mitochondrial horizontal transfer (HT) events (BTN1-HT1 to -HT6 and BTN2-HT1 to -HT3), although it is currently impossible to ascertain whether two of these lineages represent the original mtDNA haplotypes of the CedBTN founder individuals. The correspondence between mtDNA and nuclear lineages was supported by a phylogenetic tree inferred from the genotypes of nuclear BTN-specific SNVs across the entire set of 61 sequenced tumours (**Fig. S10**). Furthermore, tumours from distinct mtDNA lineages within the same CedBTN clone presented no evident cytohistological differences (**Fig. S11; Table S10**). We evaluated the potentially independent origins of the nine mtDNA lineages using three topology testing methods on the mtDNA phylogenies (Shimodaira–Hasegawa [SH] and approximately unbiased [AU] tests for the ML tree, posterior odds for the Bayesian tree), which consistently supported independent origins for eight of the lineages (*P*=0 for SH, *P*<5×10^-5^ for AU, posterior odds=0).

Analyses of the geographic distribution of mtDNA haplotypes from tumours and their sister taxa (defined as non-neoplastic samples derived from the same node in the phylogeny) provided insight into the origins and spread of CedBTN mtDNA lineages. Firstly, although most tumour samples from the same mtDNA lineage are usually found in the same geographic region (e.g. BTN1-HT1 in south Portugal, BTN1-HT2 in France, BTN1-HT3 in Ireland), this is not the case for BTN2-HT2, for which tumour specimens were collected in north-west Spain and Wales (**Fig. 2C**). Secondly, the geographic ranges of tumours and their sister taxa may be expected to overlap (e.g. BTN1-HT3 and sister taxa in Ireland), or at least be proximate (e.g. BTN1-HT2 in France and sister taxa in Spain and Portugal), yet we observed four mtDNA lineages (BTN1-HT1, BTN1-HT2, BTN2-HT2, BTN2-HT3) occupying regions distant from the ranges of their sister taxa (**Fig. 2D**). Two remarkable cases are BTN1-HT1 and BTN2-HT3, for which tumours were found in Portugal and Spain, respectively, while their sister taxa were sampled in Ireland, Germany, Denmark and Norway. Thirdly, the sister taxa of CedBTN2 mtDNA lineages were almost invariably found in northern regions (Denmark, Germany, Norway and the Netherlands), despite the fact that no CedBTN2 tumours were observed in this range (**Fig. 2D; Fig. S12**). Although we cannot rule out an anthropogenic cause for some of these patterns, the geographic structure of the mtDNA phylogeny suggests that CedBTN clones have spread over long distances along the Atlantic coast of Europe, possibly through a gradual process of natural colonization. Host mitochondria have been captured by CedBTN cells at different points in this process, potentially to replace heavily mutated incumbent mtDNA (*26, 28*).

In addition to mtDNA SNVs, inspection of sequencing depths revealed three independent amplifications spanning the control region of the mtDNA D-loop in CedBTN1, which are absent from healthy cockles (**Fig. S13**). The amplified sequences share a common start motif and overlapping microhomology at the boundaries, which is associated with imperfect DNA break repair (*29*). The evolutionary significance of these recurrent amplifications is unclear: they may be neutral changes, or the result of selfish selection at the mitochondrial level (*27*), or yet confer an advantageous phenotype on BTN cells. Notably, similar D-loop amplifications have been identified in both BTN and nonneoplastic samples from North American soft-shell clams (*30*), as well as human cancers (*31*).

Analysis of changes in mtDNA VAF across different tissues of the same animal revealed three cases in which two CedBTN mtDNA lineages coexisted within the same host (**Fig. S14**). In one remarkable animal (EICE18/910), VAF analysis revealed the presence of mtDNA haplotypes from both CedBTN1 and CedBTN2 lineages (**Fig. 2E**), with co-infection by both clones being confirmed through histopathological identification of cell morphologies matching DN types A and B (**Fig. 2F**). Histopathological re-evaluation of our tumour collection uncovered seven additional cases of coinfection by both types of DN (**Table S2**). These findings suggest that, despite its extreme rarity in mammalian transmissible cancers, host co-infection is a relatively frequent event in cockle BTN.

### Lineage-specific mutational processes operate in cockle BTN

To investigate the processes of DNA damage and repair causing mutations in CedBTN, we examined patterns of SNVs and indels at particular sequence contexts, termed mutational signatures (*32*). The mutational spectra of germline cockle SNVs and BTN-specific SNVs are broadly similar, the major difference being a higher fraction of cytosine-to-thymine (C>T) substitutions at non-CpG sites in CedBTN relative to the germ line (**Fig. 3A**). We assessed mutational processes across the CedBTN phylogeny by defining six subsets of BTN-specific variants (**Fig. 3B**): SNVs shared by all samples from each clone, but not shared between clones (two pre-divergence sets, ‘A1’ and ‘A2’; **Fig. 1D**); SNVs shared by only some tumours in each clone (two non-private post-divergence sets); and SNVs present in one tumour (two private sets). We also defined two germline sets: ancestral SNVs shared by both CedBTN clones (ancestral set, ‘A0’), and SNVs identified in three non-neoplastic cockles. While the two pre-divergence sets, containing mostly germline variants, present similar mutational spectra, the largely somatic post-divergence sets exhibit notable differences, particularly in the C>T component (**Fig. 3B**).

**Figure 3.**
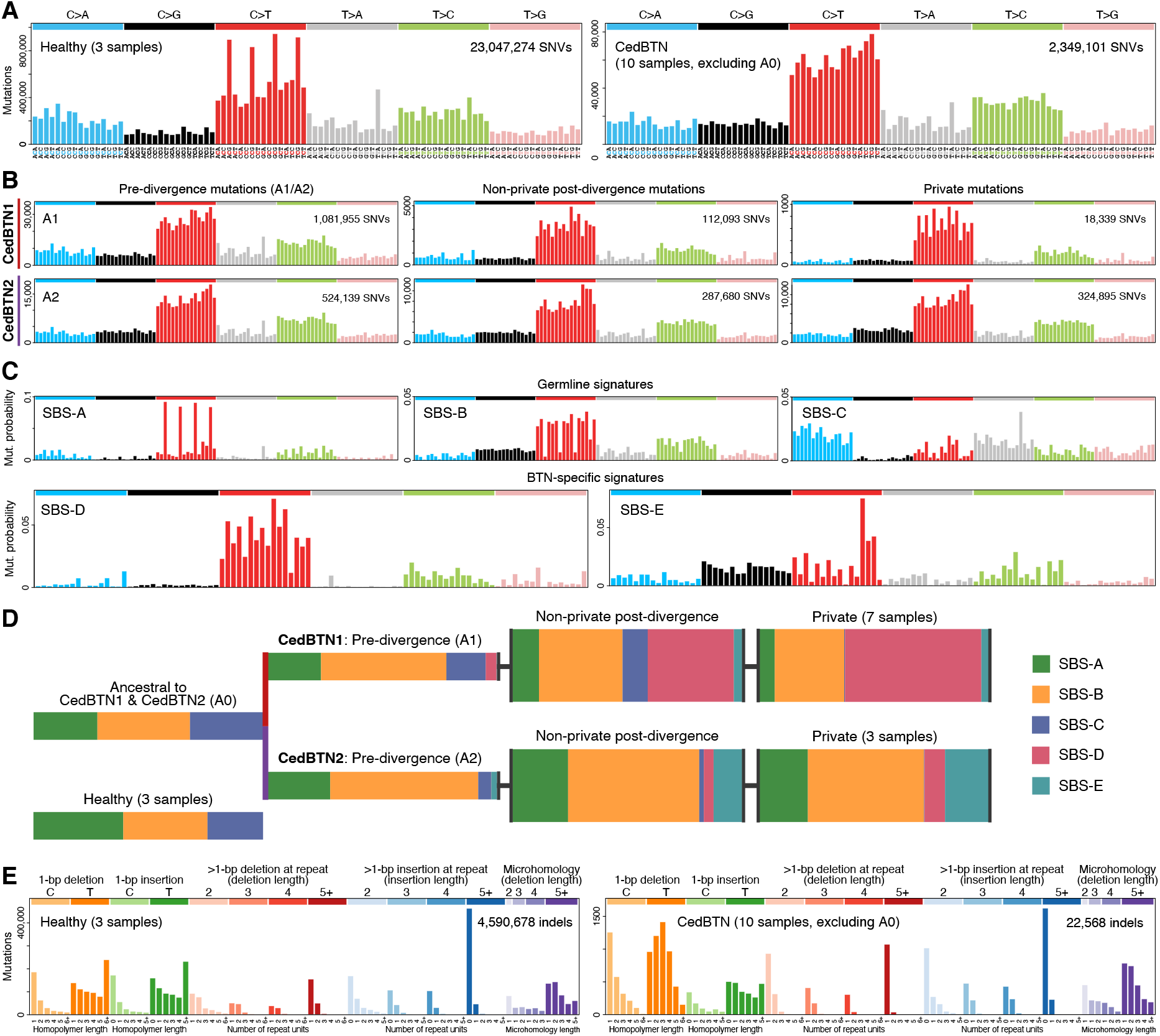
Mutational processes in CedBTN. **A**, Mutational spectra of germline SNVs in three healthy cockle samples (left) and BTN-specific SNVs in 10 CedBTN samples (excluding the set of shared ancestral SNVs, A0 in **Fig. 1D**). The *x*-axis presents 96 mutation types in a trinucleotide context, coloured by base substitution type (*29*). **B**, Mutational spectra of subsets of BTN-specific variants in CedBTN1 (top) and CedBTN2, including predivergence variants (left; A1/A2), non-private post-divergence variants (centre) and private variants. The *y*-axis presents number of mutations. **C**, Germline (top) and BTN-specific mutational signatures inferred from the spectra shown in **A** and **B** (plus the A0 spectrum), and normalized to correct for the cockle genome trinucleotide frequencies. The *y*-axis presents mutation probability. **D**, Contribution of each mutational signature to the SNVs in each segment of the CedBTN phylogenetic tree (**Fig. 1D**) and in healthy samples. Bars for post-divergence variant sets are depicted with greater width to denote collapsing of multiple internal branches of the tree. **E**, Mutational spectra of germline indels in three healthy samples (left) and BTN-specific indels in 10 CedBTN samples (excluding the shared ancestral set, A0). The *x*-axis presents 83 insertion/deletion types coloured by type and length (*29*).

With the aim of quantifying the contribution of different mutational processes to these variant sets, we applied a Bayesian approach to infer five mutational signatures *de novo* from their mutational spectra (*33*) (**Fig. 3C**). Three of these signatures (SBS-A, SBS-B, SBS-C) are shared by germline and BTN-specific sets, while the remaining two (SBS-D, SBS-E) are BTN-specific. Most signatures show similarity to human mutational signatures, especially if the latter are corrected for the trinucleotide composition of the human genome. Among the germline signatures, SBS-A probably corresponds to a mixture of human signatures SBS1 (cosine similarity 0.84), caused by spontaneous deamination of 5-methylcytosine at CpG sites (*29, 34*), and SBS5 (0.90), thought to arise from multiple endogenous mutational processes (*29, 35, 36*); SBS-B resembles human SBS40 (0.79), possibly caused by the same endogenous processes as SBS5 (*35, 36*); and SBS-C is similar to SBS8 (0.82), a signature associated with DNA repair and replication errors in human cancers and absent from the human germ line (*37, 38*). Of the BTN-specific signatures, SBS-D resembles both SBS23 (0.86), a signature of unknown aetiology described in human myeloid and brain tumours (*29*), and SBS11 (0.81), associated with the alkylating chemotherapeutic agent temozolomide (*32*); the profile of SBS-E has no evident human counterpart, the closest match being SBS40 (0.71).

To explore variation in the activity of mutational processes, we assessed mutational signature exposures across the BTN phylogeny. Signatures SBS-D and SBS-E, while undetectable in germline variant sets, are each predominantly associated with one BTN clone: whereas SBS-D dominates the spectrum of CedBTN1 post-divergence mutations, SBS-E is mainly active in the CedBTN2 post-divergence set (**Fig. 3D; Table S11**). We note that, while BTN-specific variant sets (including A0) present lower SBS-A exposures relative to the cockle germ line, this may simply reflect disproportionate filtering of variants at CpG sites, which are underrepresented relative to other sequence contexts in the cockle genome.

Inspection of indel spectra provided evidence for a variety of mutational processes in germline and BTN-specific sets (**Fig. 3E**). Although not every observed pattern can be matched to a human signature, germline indels appear to be enriched in signatures ID1 and ID2 (single-nucleotide deletions and insertions at A/T homopolymers, caused by strand slippage during DNA replication (*29*)), as well as ID9 and ID14 (single-nucleotide deletions and insertions at homopolymers, of unknown aetiology).

BTN-specific indels present lower contributions from ID1 and ID2 relative to the germ line, and seem enriched in ID5 (single-nucleotide deletions at A/T homopolymers, of unknown aetiology) and ID8 (long insertions and deletions, possibly caused by repair of DNA double-strand breaks via non-homologous end-joining (*29*)). Hence, mutational processes absent from the germ line, and possibly linked to genomic instability, appear to have contributed substantial fractions of indels to CedBTN genomes.

### Pervasive genomic instability drives the evolution of cockle BTN

Previous cellular studies have shown that cockle DN is distinguished by an unusual, broad continuum of ploidy ranging from 1.3*n* to 9.6*n*, and a variable karyotype marked by abundant microchromosomes (*39–41*). To further investigate this hallmark of DN in cockle BTN, we performed cytogenetic analysis of 261 metaphase spreads from neoplastic cells in six tumours, three from each CedBTN lineage (**Fig. S15**). This revealed extensive variation in chromosome number and size across tumours, with the median chromosome number per sample varying between 98 and 276 (**Table S12**). Notably, we also observed wide variability in chromosome number within individual tumours. For instance, neoplastic metaphase spreads from sample PACE17/478H contained 11–354 chromosomes of variable size and structure. Fluorescent *in situ* hybridization (FISH) probes targeting telomeric sequences showed that, despite such karyotypic plasticity, all the chromosomes in CedBTN cells present a canonical structure (**Fig. 4A**). These results suggest that the shifting karyotypes of CedBTN are probably the outcome of extensive chromosomal reorganization and frequent chromosome mis-segregation during anaphase. It therefore seems likely that a fraction of cell divisions may produce inviable cells in these tumours.

**Figure 4.**
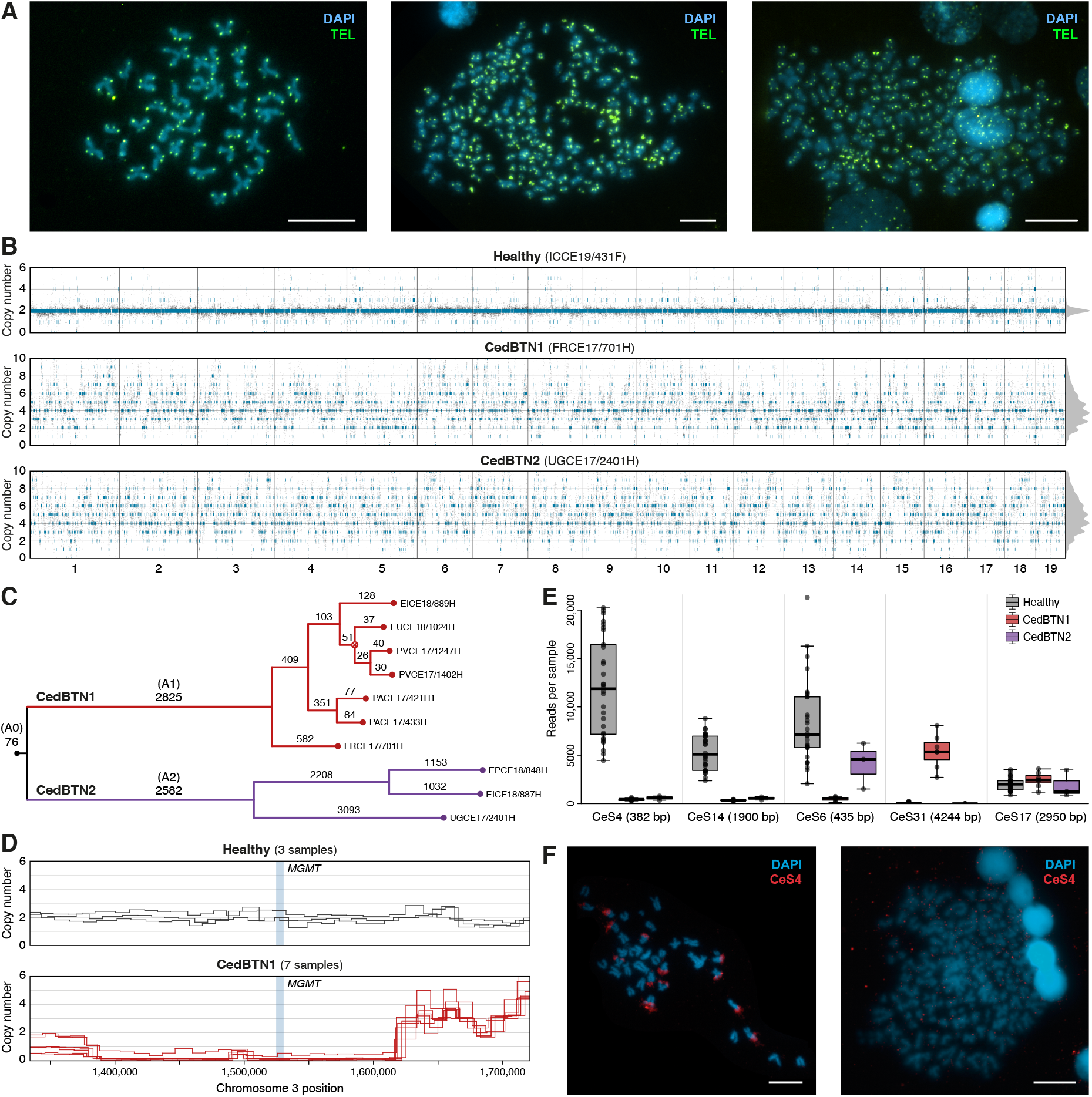
Chromosomal, copy number and structural variation in cockle BTN. **A**, FISH of telomeric peptide nucleic acid probes (TEL, shown in green) onto healthy (left) and representative type A (centre) and type B neoplastic metaphase spreads. All chromosomes, including the smallest neoplastic chromosomes, hold telomeric signals on all chromatid ends. Scale bars, 10 μm. **B**, CN profiles of representative healthy, CedBTN1 and CedBTN2 samples. Grey dots represent estimates of unrounded CN for 10-kb windows along the reference genome (*x*-axis); blue segments indicate inferred segments of integer CN. Distributions of unrounded CN are shown on the right margin. **C**, Phylogenetic tree inferred from BTN-specific SVs. The number of SVs per branch is indicated, and branches corresponding to sets of ancestral or pre-divergence variants (A0, A1, A2) are labelled. Bootstrap support values (*n*=1000 replicates) are ≥99.9 for all nodes except that marked with symbol Ä (91.6). **D**, Copy number profiles in a 500-kb region around the *MGMT* gene locus in healthy (*n*=3) and CedBTN1 (n=7) samples. Each sample is represented by a line. The highest CedBTN1 CN estimate at the gene locus (CN=0.4) corresponds to sample EICE18/889H. **E**, Numbers of sequence reads aligning to five satellite DNA elements identified in a diverse set of healthy cockles (*n*=30) and CedBTN1 (*n*=7) and CedBTN2 (*n*=3) tumours. Each dot represents a sample. Boxes represent first and third quartiles; middle line within each box denotes the median; whiskers indicate values within 1.5× interquartile range from the first and third quartiles. Monomer size is provided for each satellite. **F**, FISH of DNA probe for satellite CeS4 (red) onto representative metaphases of healthy (left) and neoplastic specimens. Scale bars, 10 μm; DAPI, 4’,6-diamidino-2-phenylindole.

Next, we inferred copy number (CN) profiles from whole-genome sequencing data for each tumour in our ‘golden set’. The profiles were marked by a ubiquitous pattern of highly complex CN alterations along every reference chromosome, with lower CN levels visibly underrepresented (**Fig. 4B**). CN distributions were consistent with a modal CN of 4*n*, suggestive of ancestral tetraploidy, except for one tumour (UGCE17/2401H) presenting a modal CN of 5*n*. Profiles were loosely conserved across tumours from each lineage, with a combination of shared and sample-specific CN features (**Fig. S16**). Moreover, CN distributions revealed a strong aberrant background of chromosomal regions with additional CN states, which in some cases obscured the expected tetramodal or pentamodal CN profile (**Fig. S16**). The cytogenetic findings above suggest that this aberrant CN background may result from persistent chromosome mis-segregation, possibly resulting in intra-tumour heterogeneity in CN. Such heterogeneity is probably amplified by cell transmission bottlenecks to produce the observed intertumour CN variability. Overall, our analyses indicate that both CedBTN clones are highly aneuploid, polyploid lineages that suffered at least one whole-genome duplication event in early tumorigenesis, leading to a likely tetraploid state that, in the case of CedBTN2, later developed further CN gains in the UGCE17/2401H branch.

To further characterize the landscape of somatic alterations in cockle BTN, we applied multiple established algorithms to call structural variants in the 10 ‘golden set’ tumours. We then removed potentially germline events by genotyping these variants on 455 non-neoplastic samples. This approach yielded a conservative set 18,272 high-confidence structural variants (6,916 in CedBTN1, 10,925 in CedBTN2), with deletions being the most frequent type of event (80%, 14,589/18,272; **Fig. S17**). A maximum-parsimony phylogenetic tree reconstructed from these variants confirmed the CedBTN nuclear phylogeny inferred from SNVs, supporting two divergent lineages with a minimal fraction of shared structural variants (**Fig. 4C**).

Although analysis of dN/dS ratios revealed no evidence of positive selection for post-divergence SNVs or indels in either clone, the availability of CN data offered an additional opportunity to identify potential early cancer-driver alterations. We systematically screened for gains (CN≥8) and losses (CN≤1) of regions containing oncogenes and tumour suppressor genes, respectively. This analysis detected likely ancestral amplification of two canonical oncogenes in CedBTN1: *MDM2* (10–13 copies in CedBTN1; mean CN=10.9; gene CN percentile=98.3), encoding an important inhibitor of tumour suppressor proteins, and *CCND3* (8–18 copies in CedBTN1; mean CN=10.7; gene CN percentile=98.2), encoding a cyclin that promotes the G1/S cell cycle transition (**Table S13**). Recurrent amplification of these genes has been observed in multiple cancer types, and is thought to prevent cell cycle arrest and apoptosis under conditions of genomic instability (*42–45*). Notably, we also identified an ancestral homozygous deletion of *MGMT* in CedBTN1 (**Fig. 4D; Table S13)**. The enzyme encoded by this gene, *O*^6^-methylguanine-DNA methyltransferase, is essential for repair of alkylated DNA bases, and its inactivation results in hypersensitivity to the toxic and mutagenic effects of alkylating agents (*46–48*). Given the cumulative and virtually lineage-specific activity of signature SBS-D (**Fig. 3D**), and its similarity to COSMIC SBS11 (caused by the alkylating agent temozolomide (*32*)), SBS-D most likely reflects unrepaired alkylation of DNA bases due to loss of *MGMT*. The resemblance between SBS-D and SBS23 further suggests that SBS23 may arise from deficient DNA alkylation repair in human cancers.

### Satellite DNA expansions illuminate the emergence of CedBTN

Finally, we applied a computational method to examine the repetitive complement of the *C. edule* genome, with a focus on satellite DNA. These repetitive sequences are relevant for genome stability, exhibiting long-term conservation and propensity for rapid copy number changes (*49*). Our method identified 34 satellite DNA candidates in the common cockle reference genome (**Table S14**), four of which varied in frequency between non-neoplastic and BTN genomes, providing further insights into the origins of cockle BTN (**Fig. 4E**). Two satellites, named CeS4 and CeS14, were found at high frequency in all samples from a genetically diverse cohort of non-neoplastic cockles, yet were entirely absent from both BTN clones. We designed FISH probes to target satellite CeS4, which confirmed the results obtained from sequencing data (**Fig. 4F**). This finding suggests that both CedBTN1 and CedBTN2 may be ancient cancer lineages that diverged from the cockle population before the emergence and expansion of CeS4 and CeS14 in the *C. edule* germ line. Another satellite, CeS6, was found in cockle populations and CedBTN2 samples, while being absent from CedBTN1 (**Fig. 4E**). Lastly, despite satellite CeS31 being exclusive to CedBTN1, our data did not support exclusive presence of any satellite DNA in CedBTN2 samples. These observations suggest that CedBTN2 may have diverged from the cockle population more recently than CedBTN1.

## Discussion

Despite several BTN clones having been newly described in recent years (*9–14*), no analyses of whole BTN genomes have yet been reported. Combining a range of approaches, our study provides a first outlook into the genomes of these singular marine leukaemias in European common cockles, complementing the work of Hart *et al. (30*) on American soft-shell clams. Both studies reveal neoplastic genomes marked by structural instability. In the case of cockle BTN, we find evidence for ongoing extreme genomic instability, most likely activated by early whole-genome duplication (*50*) and fuelled by recurrent chromosome mis-segregation during mitosis (*51*). This is in stark contrast with the three transmissible cancers described in terrestrial mammals (dogs and Tasmanian devils), which present remarkable karyotypic stability (*1, 5, 52*), and thus challenges the notion that development of a durable genome architecture is required for long-term survival of cancer lineages. Although our data do not allow estimation of precise ages for cockle BTN, multiple lines of evidence suggest that these clones may have emerged centuries or millennia ago. These include the broad geographic distribution of tumours, the marked genetic divergence between tumours and modern cockles, the recurrent capture of host mitochondria by tumours, and the absence in tumours of satellite DNA elements that are vastly expanded in the cockle germ line. Furthermore, Hart *et al*. estimate an age of ~500 years for the BTN clone affecting soft-shell clams (*30*), supporting that long-term survival of marine transmissible cancers is possible. Taken together, our findings suggest that CedBTN lineages have undergone a relatively long history of pervasive genomic instability. Studying the mechanisms that enable BTN cells to overcome the effects of such instability will broaden our understanding of the conditions required for cancers to survive and adapt over the long term.

## Acknowledgements

We thank the fishermen’s associations (‘cofradías’) of Galicia for their advice and assistance with sampling, and the Galicia Supercomputing Centre (CESGA) for the availability of informatic resources. We thank L.F. Møller (National Institute of Aquatic Resources, Denmark), B. Hussel (Alfred Wegener Institute), M. Wolowicz (University of Gdansk), T. Verstraeten (Ghent University), M.L. Martínez and A.M. Insua Pombo (Universidade da Coruña), C. García de Leaniz (Swansea University), A. Smith and A. Harvey (Marine Biological Association, UK), R. Parks (Centre for Environment, Fisheries and Aquaculture Science, UK) and T. Magnesen (Universitetet i Bergen) for providing samples for this project. We thank C. Canchaya (Universidade de Vigo), M. Rey, J. Quinteiro, M. Hermida and P. Martínez (Universidade de Santiago de Compostela) for helpful advice on genome assembly and annotation, M. Rodríguez (Universidade de Vigo) for administrative support, and E.P. Murchison (University of Cambridge) for helpful advice and critical reading of the manuscript.

## Author contributions

A. Villal., D.P. and J.M.C.T. designed the project. A.L.B., M.Sa., D.G.-S., S.D., S.R., J.Z., M.A.Q., I.O., J.J.P., J.D. and A.B.-O. developed methods. A.L.B., M.Sa., D.G.-S., S.D., S.R., J.Z., Y.L., I.O., J.T., Y.S.J., J.D. and A.B.-O. performed computational analyses. T.P., L.T., J.A., Z.N. and D.P. assisted with analyses. A.L.B., M.Sa., D.G.-S., S.D., A. Villan., A.P.-V, A.V.-F., J.T., J.R.-C., P.A., J.A. and J.J.P. performed laboratory work. A.L.B., D.G.-S., S.D., M.A.Q., A.P.-V, J.T. and J.R.-C. performed sequencing methods. A.L.B., D.G.-S., S.D., A.V.-F., A. Villan., D.C., R.R., J.A., A.M.A, P.B., R.C., B.E.K., U.I., X.M., N. G.P., I.P., F.R., P.R., M.Sk. and K.S. provided samples. A.L.B., M.Sa., D.G.-S., S.D., S.R., J.Z., Y.L., J.J.P., Y.S.J., D.P., J.D., A.B.-O. helped with interpretation of results. A.L.B., D.G.-S., S.D., A.P.-V, J.R.-C., A. Villan., P.A., J.A. performed sample management. A.C., D.I., M.J.C, A. Villal., Z.N. and D.P. provided technical advice. A.L.B., M.Sa., D.G.-S., S.D. Y.L., J.Z., J.D. and A.B.-O. generated figures. A.B.-O. and J.M.C.T. wrote the manuscript with contributions from all other authors.

